# Actin dynamics regulation by TTC7A/PI4KIIIα axis limits DNA damage and cell death during leukocyte migration

**DOI:** 10.1101/2021.10.14.464382

**Authors:** Tania Gajardo, Marie Lô, Mathilde Bernard, Claire Leveau, Marie-Thérèse El-Daher, Mathieu Kurowska, Gregoire Le Lay, Despina Moshous, Bénédicte Neven, Alain Fischer, Gaël Ménasché, Geneviève de Saint Basile, Pablo Vargas, Fernando E. Sepulveda

**Affiliations:** 1Molecular basis of altered immune homeostasis laboratory, INSERM UMR 1163, Paris, France; Université de Paris, Imagine Institute, F-75015, Paris France; UMR 144, Institut Curie, Paris, France; Institut Pierre-Gilles de Gennes, PSL Research University, Paris, France; Pediatric Immunology, Hematology and Rheumatology Department, Necker-Enfants Malades University Hospital, Assistance Publique Hôpitaux de Paris, Paris, France; Collège de France; Centre d’Etude des Déficits Immunitaires, AP-HP, Hôpital Necker-Enfants Malades, Paris, France; Centre national de la recherche scientifique – CNRS

**Keywords:** Cell migration, leukocytes, TTC7A, actin cytoskeleton, DNA damage, cell death.

## Abstract

The actin cytoskeleton has a crucial role in the maintenance of the immune homeostasis by controlling various cell processes, including cell migration. Mutations in the *TTC7A* gene have been described as the cause of a primary immunodeficiency associated to different degrees of gut involvement and alterations in the actin cytoskeleton dynamics. Although several cellular functions have been associated with TTC7A, the role of the protein in the maintenance of the immune homeostasis is still poorly understood. Here we leverage microfabricated devices to investigate the impact of TTC7A deficiency in leukocytes migration at the single cell level. We show that TTC7A-deficient leukocytes exhibit an altered cell migration and reduced capacity to deform through narrow gaps. Mechanistically, TTC7A-deficient phenotype resulted from impaired phosphoinositides signaling, leading to the downregulation of the PI3K/AKT/RHOA regulatory axis and imbalanced actin cytoskeleton dynamic. This resulted in impaired cell motility, accumulation of DNA damage and increased cell death during chemotaxis in dense 3D gels. Our results highlight a novel role of TTC7A as a critical regulator of leukocyte migration. Impairment of this cellular function is likely to contribute to pathophysiology underlying progressive immunodeficiency in patients.

---

Autosomal recessive biallelic mutations in the gene coding for the tetratricopeptide repeat Domain 7A (*TTC7A*) have been identified as the cause of an immune and gastrointestinal disorder of variable severity ^1–4^. Depending on the type of mutation, gastrointestinal symptoms can present as very-early-onset inflammatory bowel disease (VEO-IBD) or multiple intestinal atresia (MIA). On the other hand, immune manifestations of TTC7A-deficient patients range from mild lymphopenia to combined immunodeficiency ^5^. In general, TTC7A-deficient patients develop a progressive leukopenia, leading to increased susceptibility to infections.

The TTC7A protein contains 9 tetratricopeptide repeat (TPR) domains, which have been proposed to act as scaffold for protein complexes^6^. Our group and others have described diverse functions mediated by TTC7A. In vitro, cells from TTC7A-deficient patients present with disrupted actin cytoskeleton and cell polarity, through the increase of RhoA-mediated signaling ^1^. TTC7A also interacts with the supramolecular complex containing phosphatidylinositol 4 kinase type III α (PI4KIIIα), EFR3 homolog B and FAM126, in the plasma membrane ^7^. PI4KIIIα is required for the synthesis of phosphatidylinositol 4-phosphate (PI4P), that is necessary for plasma membrane identity, cell survival and cell polarity ^8, 9^. TTC7A can also be localized in the nucleus, participating in the regulation of chromatin structure and nuclear organization ^10^. Finally, in mice, Ttc7 controls hematopoietic stem cells stemness ^11^. Despite of our improved understanding of the different cellular functions of TTC7A, the pathophysiological mechanisms underlying TTC7A-associated immunodeficiency are still not well characterized.

In the present study, we leverage the use of microfabricated devices to investigate the impact of TTC7A deficiency on the migratory capacity of leukocyte at the single cell level. We found that TTC7A-deficient leukocytes were characterized by an increased cell speed compared to control cells but failed to deform their nucleus when migrating along micrometric spaces. Mechanistically, TTC7A deficiency disrupted actin cytoskeleton dynamics downstream the PI4KIIIα/PI3K/AKT signaling pathway. Notably, chemotactic migration of leukocytes from TTC7A-deficient patients in dense 3D microenvironments resulted in increased DNA damage and cell death. We propose that altered actin dynamics observed in TTC7A deficient leukocytes modifies their migratory capacity and survival in complex 3D microenvironments, possibly contributing to progressive leukopenia observed in TTC7A-deficient patients.

## Results

### Ttc7 regulates 1D migration of immature murine dendritic cells

We have previously shown that gut organoids derived from TTC7A-deficient patients present with an altered polarity due to increased RHOA signaling^1, 2^. This pathway has also been shown essential for leukocyte migration under confinement ^12, 13^. Hence, we hypothesized that Ttc7 deficiency could affect leukocyte migration. To test this hypothesis, in a first step we generated bone marrow-derived dendritic cells (DC) from *flaky skin* (*fsn*) mice (natural mutant deficient for *Ttc7*). We observed that Ttc7 was not required for DC differentiation, nor for DC capacity to respond to TLR stimulation in vitro, as a similar up-regulation of MHC-II and co-stimulatory molecules (CD40, CD80, CD86 and CCR7) occurred in Ttc7-deficient and control cells (fig 1a and S1a). In addition, both cell types secreted similar levels of inflammatory cytokines in response to TLR4 stimulation (Supplementary fig S1b). To assess whether Ttc7 is required for DC motility, we compared the migration of immature (iDC) and LPS-activated DC (mDC) from control and *fsn* mice. To do so, we used microchannels consisting in small tubes in which cells migrate in 1 dimension (1D) ^13^. As previously described, mature control DC (mDC^ctrl^) had an increased speed compared to control immature DC (iDC^Ctrl^) (fig 1b-c and Supplementary Video 1). Interestingly, no difference in cell speed associated to DC maturation was observed when comparing Ttc7-deficient iDC (iDC*^fsn^*) and mDC (mDC*^fsn^*), since both populations had a similar high migration speed, resembling the behavior of mDC^Ctrl^ (fig 1b-c and Supplementary Video 1). Increased cell speed of iDC*^fsn^* was observed for different levels of confinements (i.e., using microchannels of 4 and 8µm width, respectively) (supplementary fig S1c). Similarly, iDC*^fsn^* behavior was independent of cell adhesion and autocrine sensing of chemokines, since equivalent results were observed when using polyethylene glycol (PEG) coated microchannels (to block integrins interaction with the surface) and in cells treated with pertussis toxin (PTX – to block G protein-coupled chemokine receptors) (supplementary fig S1d-e)^14^. In addition, the impact of Ttc7 in the control of iDC speed was likely independent of genetic background, as comparable results were obtained in control and *fsn* DC obtained from Balb/c mice (supplementary fig S1F). Ttc7 also controlled migration of T cells as similar results were obtained when comparing control and Ttc7-deficient T cells (supplementary fig S1g). Altogether, we concluded that Ttc7 is a critical regulator of leukocyte migration under confinement *in vitro*.

**Figure 1.**
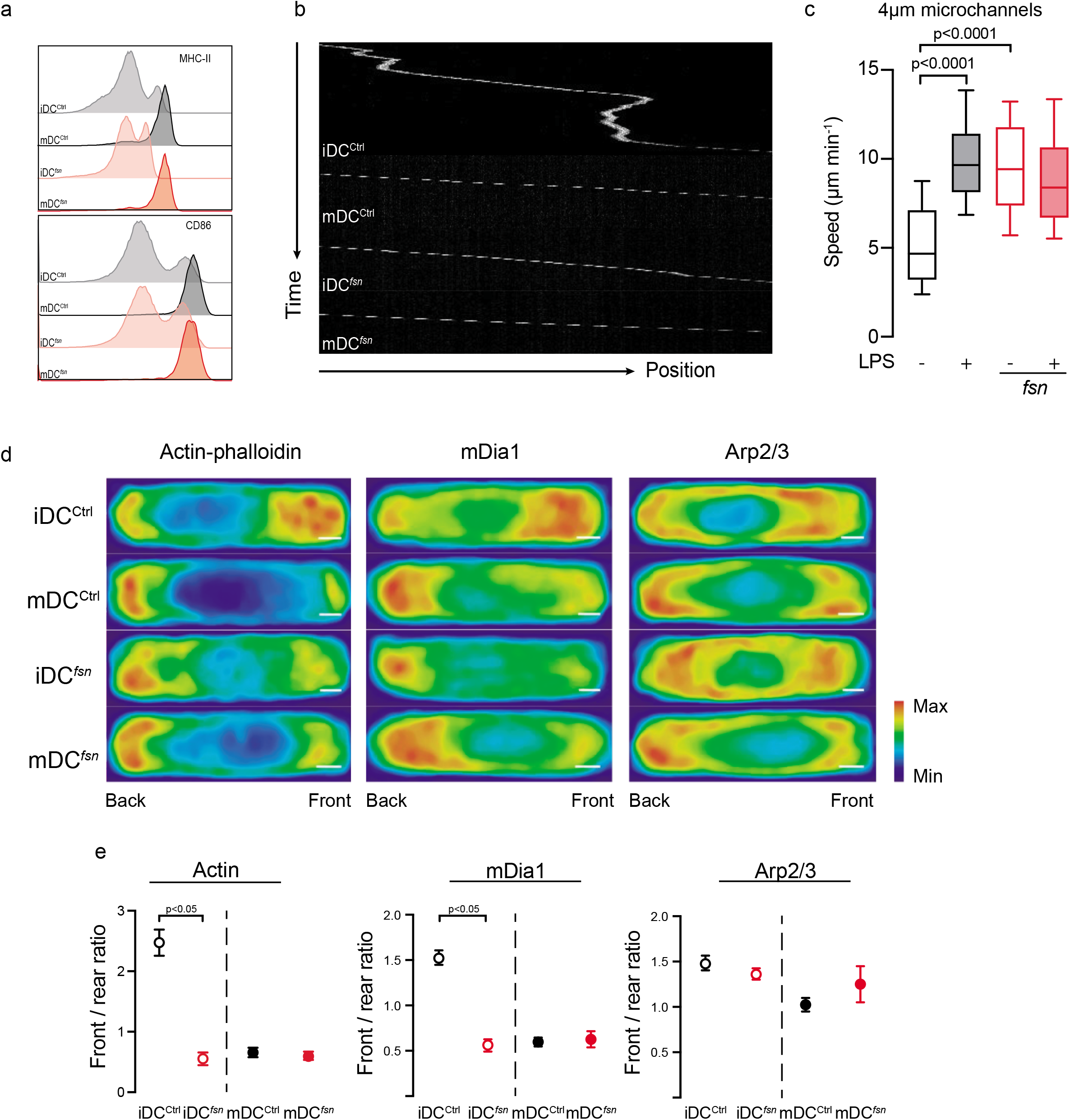
Ttc7 is necessary for 1D migration of DCs: Analysis of DC^ctrl^ and DC*^fsn^* migrating in 4µm x 5µm fibronectin-coated microchannels. Cell imaging was performed between 4 and 16 hours after LPS stimulation (100ng/ml, 30 min) or not. (a) Flow cytometry analysis of MHC-II and CD86 iDC^ctrl^ (Light grey), mDC^ctrl^ (Dark grey), iDC*^fsn^* (Light red) and mDC*^fsn^* (Dark red). (b) Representative kymographs of iDC^ctrl^ (top panel), mDC^ctrl^ (upper middle panel), iDC*^fsn^* (lower middle panel) and mDC*^fsn^* (bottom panel), migrating in microchannels. (c) Mean instantaneous speed of iDC^ctrl^ (Black-empty bars), mDC^ctrl^ (Black-filled bars), iDC*^fsn^* (Red-empty bars) and mDC*^fsn^* (Red-filled bars). One representative experiment of 10 is presented. Boxes include the 80% of the points; bars represent the higher and lower 10% of points. One-way ANOVA test was used to evaluate statistical significance. (d) Density maps of iDC^ctrl^ (top panels), mDC^ctrl^ (upper middle panels), iDC*^fsn^* (lower middle panels) and mDC*^fsn^* (bottom panels) fixed while migrating and stained with actin-phalloidin (right panel), mDia1 (middle panel) and Arp2/3 (left panel). One representative experiment out of 3 is presented. Scale bar 2µm. (e) Quantification of signal intensity showed as front / rear ratio for iDC^ctrl^ (Black-empty point), iDC*^fsn^* (Red-empty point), mDC^Ctrl^ (Black-filled point) and mDC*^fsn^* (Red-filled point) for the density maps in d. Actin-phalloidin (left panel), mDia1 (middle panel) and Arp2/3 (right panel). T-test was used to evaluate statistical significance.

Recently, we showed that DC migration in microchannels is controlled by actin nucleation ^14, 15^. In iDC, actin filaments are in constant oscillation between the cell front and rear. During slow motility phases, the Arp2/3 complex promotes actin nucleation at the cell front. Conversely, during fast migration phases, the mDia1 nucleator promotes F-actin polymerization at the cell’s rear. In LPS-activated DC, actin mostly relies on mDia1 function, resulting in an increased velocity during migration ^14, 15^. Therefore, we sought to determine whether cell speed is increased in absence of Ttc7 altered actin cytoskeleton dynamics. To test this, we bred LifeAct-GFP mice ^16^ with *fsn* mice to visualize the actin cytoskeleton in Ttc7-deficient cells. As expected, iDC^ctrl^ presented with a bimodal concentration of F-actin at the cell front and rear (fig 1d), while mDC^Ctrl^ presented F-actin structures preferentially at the back of the cell (fig 1d). In agreement with the fast migration speed, both iDC*^fsn^* and mDC*^fsn^* presented with F-actin preferentially concentrated at the cell rear (fig 1d-e). Further analysis of the sub-cellular distribution of mDia1 and Arp2/3, showed that mDia1 distribution followed F-actin in all conditions, whereas Arp2/3 was unchanged between control and Ttc7-deficient cells (fig 1d-e). Since mDia1 and Arp2/3 protein levels were similar between the conditions (supplementary fig S1h), these data indicate that iDC*^fsn^* actively relocates F-actin and mDia1 at the cell rear.

Collectively these results revealed that Ttc7 deficiency alters leukocyte migration by affecting the subcellular distribution of the actin nucleator mDia1 and the actin dynamics in DC.

### Increased RhoA/mDia1 activity is responsible of increased cell speed of iDC^fsn^

We and others have previously shown that gut organoids cells from TTC7A-deficient patients display an elevated RhoA/Rock-mediated signaling, resulting in altered actin cytoskeleton polarity ^2^. Therefore, we sought to determine the contribution of Rho-associated protein kinase (Rock) and myosin II to iDC motility. To do so, iDC^Ctrl^ and iDC*^fsn^* were treated with Rock inhibitor Y-27632 or myosin II inhibitor blebbistatin, and cell speed was assessed in microchannels. Both inhibitors decreased DC speed in control and Ttc7-deficient cells (fig 2a-b). RhoA signaling leads to both Rock and mDia1 activation^17^. To assess the contribution of the formin mDia1 to iDC*^fsn^* phenotype, we treated control and Ttc7-deficient iDC with the formin inhibitor Smifh2. While inhibition of mDia1 did not affect iDC^ctrl^ speed, this reduced iDC*^fsn^* speed back to control level (fig 2c). Speed normalization of iDC*^fsn^* correlated with a switch in F-actin structures from the rear to the front of the cell, recapitulating the behavior observed in control cells (fig 2d-e). In keeping with these observations, the Rock activator calpeptin specifically increased cell speed of iDC^Ctrl^, mimicking the phenotype observed in Ttc7-deficient iDC (fig 2f). In agreement with the normal cellular localization of Arp2/3 (fig 1d-e), interfering with Arp2/3 activity did not affect iDC*^fsn^* speed (fig 2g). Collectively, these data show that Ttc7 deficiency increases iDC speed under confinement by increasing F-actin polymerization at the rear of the cells in a RhoA/mDia1-mediated process.

**Figure 2.**
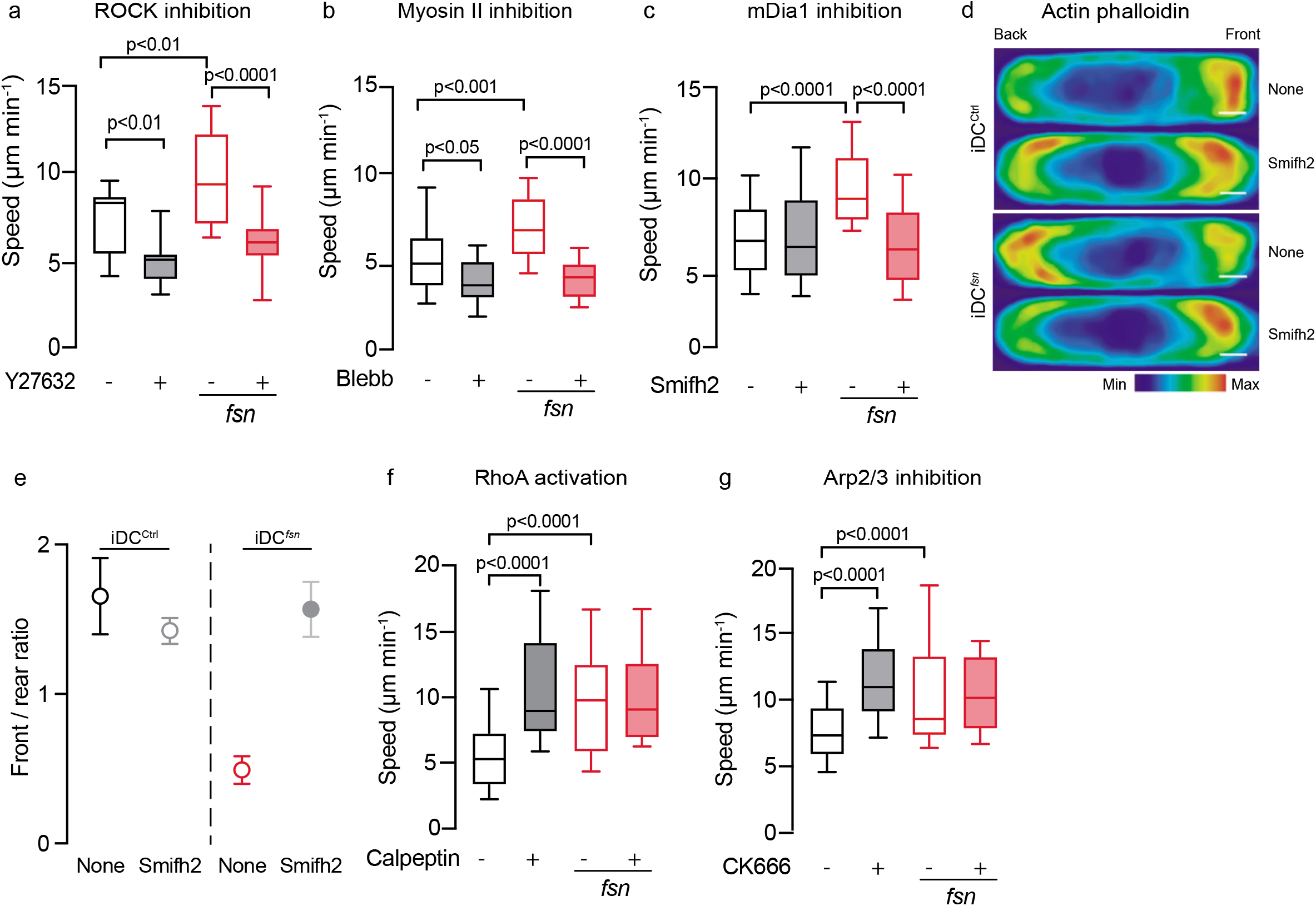
iDC^fsn^ have an increase activity of mDia1 controlling migration: Analysis of iDC^ctrl^ (Black-empty bars), treated-iDC^ctrl^ (Black-filled bars), iDC*^fsn^* (Red-empty bars) and treated-iDC*^fsn^* (Red-filled bars) migration in fibronectin-coated microchannels. Cells were incubated with the indicated drug concentration for 30 min and maintained during acquisition. Boxes include the 80% of the points and bars represent the higher and lower 10% of points. One-way ANOVA test was used to evaluate statistical significance. (a) ROCK inhibitor (Y-27632, 7.5 µM). (b) Myosin-II inhibitor (Blebbistatin, 50 µM). (c) Formins inhibitor (Smifh2, 2 µM). (d) Density maps of iDC^ctrl^ (top panels) and iDC*^fsn^* (bottom panels) treated with Smifh2 or not (“None”) as indicated, fixed while migrating and stained with actin-phalloidin (right). n≥27 cells per condition. Scale bar 2µm. (e) Quantification of signal intensity from density maps in d, for actin-phalloidin showed as front / rear ratio for iDC^ctrl^ (Black-empty point), smifh2-treated iDC^Ctrl^ (Grey-empty point), iDC*^fsn^* (Red-empty point) and smifh2-treated iDC*^fsn^* (Grey-filled point). (f) RhoA activator (Calpeptin, 30mg/ml). (g) Arp2/3 inhibitor (CK-666, 25 µM).

### TTC7A controls leukocyte migration in humans

To determine the impact of TTC7A deficiency in migration of human leukocytes, we assessed spontaneous speed of blood T cell derived from different patients carrying biallelic mutations in TTC7A (TTC7A^L304fsX59^, TTC7A^E71K^ and TTC7A^R325Q^) and healthy donors (HD) in 1D-confined microchannels. Activated control T cells had persistent trajectories (very few directional changes) and migration speed was high (fig 3a-b and Supplementary Video 2). Similar trajectories were observed with TTC7A-deficient cells. However, T cells derived from TTC7A-deficient patients, were faster than their control counterparts (fig 3a-b and Supplementary Video 2). Consistently with a general role of TTC7A in the control of leukocyte migration, similar observations were made with patient-derived lymphoblastoid B cell lines (LCL) and primary monocytes (fig 3c-d, supplementary fig S2a-b and Supplementary Video 3). In all these cases (and thus independently of the underlying mutation), TTC7A-deficient cells were faster than control cells.

**Figure 3.**
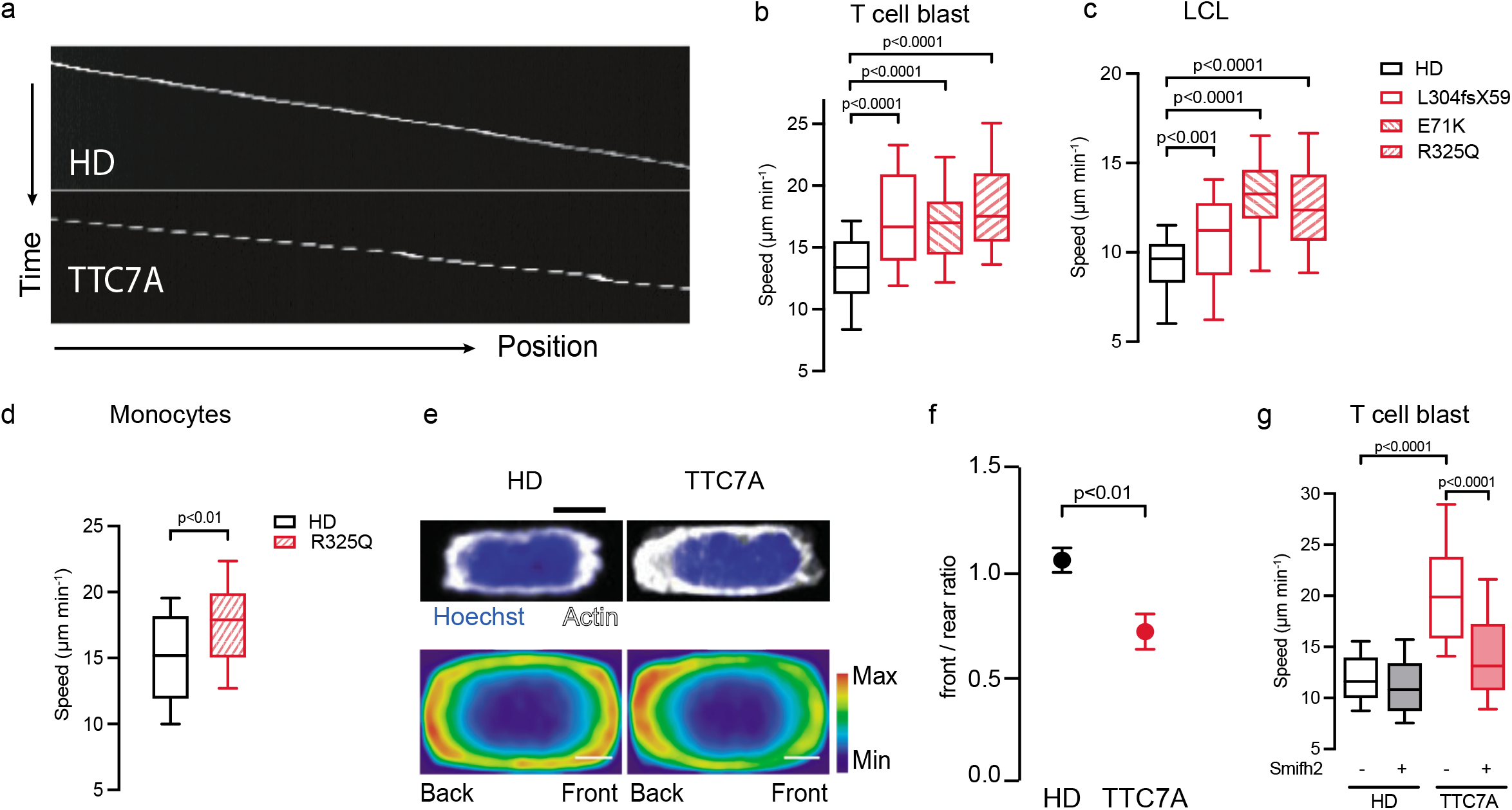
Patient derived cells recapitulate the phenotype described in mouse: Analysis of Healthy donor (HD, black-empty bars) and TTC7A (red bars) cell migration in 4µm x 5µm fibronectin-coated microchannels. Boxes include the 80% of the points and bars represent the higher and lower 10% of points. One-way ANOVA test was used to evaluate statistical significance. (a) Representative kymographs of T cell blast from HD (Top panel) and TTC7A (bottom panel). (b) Mean instantaneous speed of HD and TTC7A T cell blasts. Three mutations were analyzed, TTC7A^L304fsX59^, TTC7A^E71K^ and TTC7A^R325Q^. One experiment representative of 9 is presented. (c) Mean instantaneous speed of HD and TTC7A deficient LCL carrying one of three different mutations in the *TTC7A* gene as indicated, migration was performed in 8µm x 5µm fibronectin-coated microchannels. One experiment representative of 15 is presented. (d) Mean instantaneous speed of HD and TTC7A-derived primary monocytes. One experiment. Mann-Whitney test was used to evaluate statistical significance. (e) Representative Hoechst and actin-phalloidin immunofluorescence (top panels) from HD and TTC7A T cell blast fixed while migrating and density maps (bottom panels) generated by averaging the signal from n≥33 cell in each condition. One representative experiment out of 4 is presented. Scale bar 2µm. (f) Quantification of signal intensity of phalloidin-actin showed as front / rear ratio for HD and TTC7A T cell blast. t-test applied for statistical analysis. (g) Mean instantaneous speed of HD and TTC7A T cell blast, treated (Black and Red filled bars, respectively) or not with 2µM of Smifh2 for 30 min. One experiment representative of three is presented.

Similarly to mouse iDC, human T cells derived from healthy donors had a bimodal distribution of actin in the front and rear of the cell (fig 3e-f). In agreement with their increased cell speed, TTC7A-deficient T cells exhibited an increased F-actin polymerization at the cell rear (fig 3e-f). To evaluate the contribution of DIAPH1 (human homolog of mDia1) to this phenotype, we treated HD and TTC7A-deficient T cells with Smifh2 and measured cell migration. As shown in figure 3g, DIAPH1 inhibition restored cell speed to control values (fig 3g). These results suggest that increased migration speed under confinement of TTC7A-deficient T cells, involve an aberrant activation of DIAPH1 and F-actin polymerization at the cell rear, highlighting TTC7A as a critical regulator of actin dynamics and the migratory capacity of leukocytes as well.

### Increased cell speed observed in TTC7A-deficient cells is mediated by reduced PI4KIIIα activity

TTC7A plays a critical and highly conserved role in the assembly and function of the multi-protein complex involving PI4KIIIα, EFR3A and FAM126^18^. TTC7A deficiency leads to an impaired formation and localization of this complex, hindering PI4KIIIα activity and the subsequent phosphoinositide metabolism^3, 6^ (fig 4a). To characterize the contribution of PI4KIIIα in leukocyte migration under confinement, we treated control human T-cell blasts with two different chemical inhibitors of PI4KIIIα, BF-738735 and GSK-A1. In both cases, we observed that PI4KIIIα inhibition increased T cell speed (fig 4b), suggesting that the enhanced motility of patient-derived cells could be caused by a defective PI4KIIIα activity. To address this question, we supplemented control and TTC7A-deficient T cells with exogenous PI4P, the metabolite produced by PI4KIIIα, and assessed 1D-confined cell migration. PI4P supplementation did not affect migration of control T cells (fig 4c). In contrast, PI4P treatment reduced migration speed of TTC7A-deficient T cells and redistributed the F-actin pool towards the front of the cell (fig 4c-d). Therefore, PI4KIIIα activity modulated leukocyte speed by controlling F-actin polymerization.

**Figure 4.**
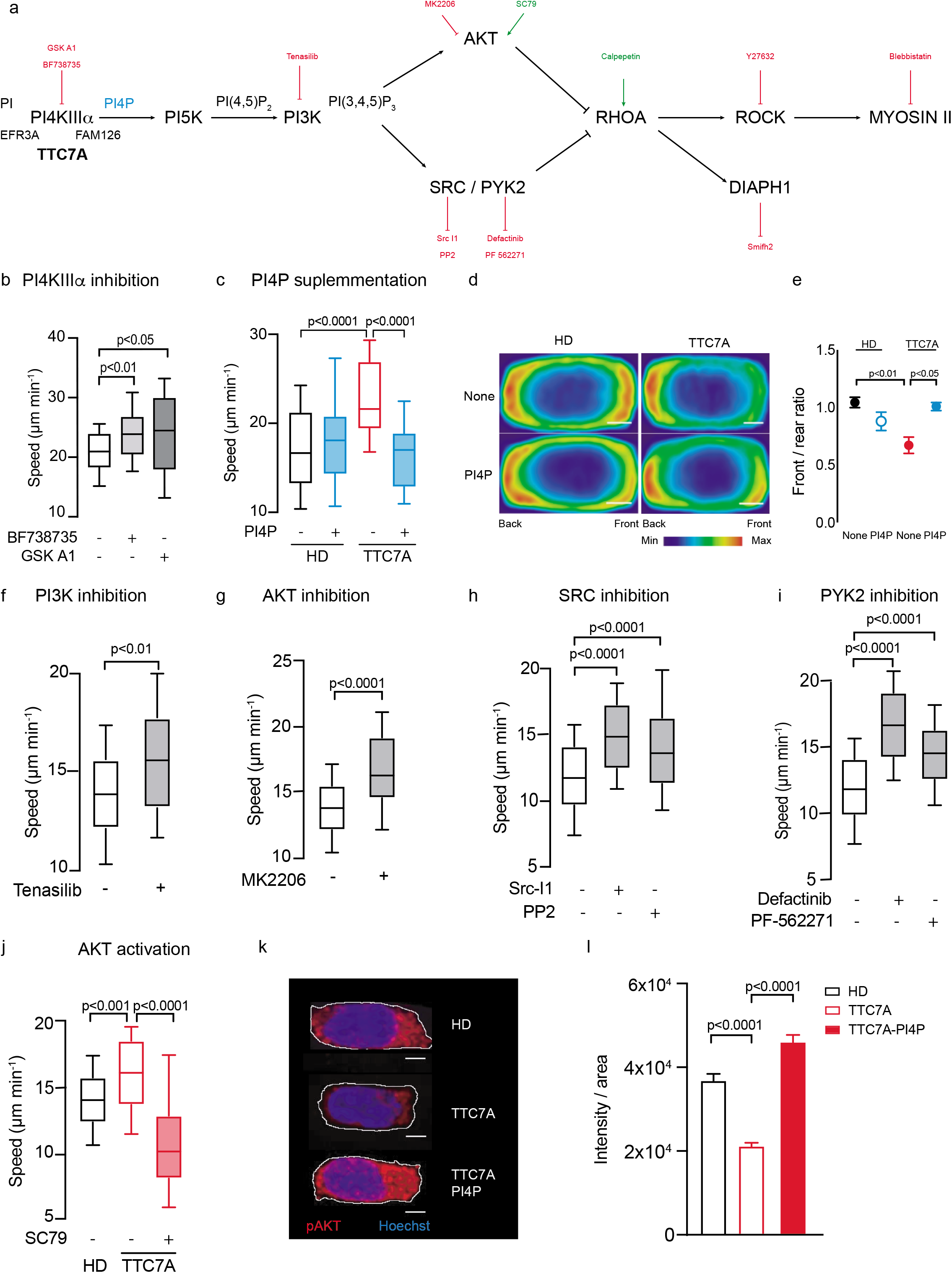
Kinase activity of PI4KIIIα controls leukocytes’ migration: Chemical intervention of HD and TTC7A T cells. (a) Schematic representation of the PI4KIIIα / TTC7A to RhoA / ROCK regulatory pathway, including the chemical modulators used in this study. Inhibitors (red), activators (green) and PI4P (blue). (b-i) Mean instantaneous speed of HD (Black-empty bar) and/or TTC7A deficient (Red-empty bar) T cell treated (Filled bars) or not with different chemical modulators. Migration was assessed in 4µm x 5µm fibronectin-coated microchannels. Boxes include the 80% of the points and bars represent the higher and lower 10% of points. One-way ANOVA or Mann-Whitney test was used to evaluate statistical significance, as appropriate. One experiment representative of three is presented in each case. (b) HD cell treated with inhibitors of PI4KIIIα. 1.7µM of BF738735 (middle) and 100 nM of GSK-A1 (right) were used. (c) HD and TTC7A cell treated (Blue-filled bars) or not with 25 µM of PI4P. (d) Density maps of HD (left panel) and TTC7A (right panel) T cells, treated (bottom panels) or not (top panels) with PI4P, fixed while migrating and stained with actin-phalloidin. n≥30 cell each condition. Scale bar 2µm. (e) Quantification of signal intensity from density maps in d, for actin-phalloidin showed as front / rear ratio for HD (Black-filled point), HD-PI4P (Blue-empty point), TTC7A (Red-filled point) and TTC7A-PI4P (Blue-filled point). T-test was used to evaluate statistical significance. (f) HD cell treated with PI3K inhibitor (Tenasilib, 45 nM). (g) HD cell treated with AKT inhibitor (MK2206-2HCl, 7.5 nM). (h) HD cell treated with inhibitors of SRC. 1µM of SRC-Inhibitor1 1µM (middle) and 10µM of PP2 (right) were used. (i) HD cell treated with inhibitors of PYK2. 1µM of Defactinib (middle) and 10µM of PF-562271 10µM (right) were used. (j) HD and TTC7A T cells treated with AKT activator SC79 (16.5 µM). (k) Immunofluorescence of T cell blast fixed while migrating and stained with pAKT and Hoechst. HD (top panel), TTC7A (middle panel), TTC7A-PI4P (bottom panel). Representative image from 2 experiments, ≥24 cells analyzed in each condition and experiment. (l) Quantification of intensity of fluorescence / area of cells in (J). HD n=62, TTC7A n= 66 and TTC7A-PI4P n=70. Fluorescence intensity determined using Icy. Scale bar 2µm.

Being the precursor of PI(4,5)P_2_ and PI(3,4,5)P_3_, PI4P availability is crucial for the phosphoinositide cascade, PI3K activity and further downstream signaling^19^. The PI3K/AKT pathway has been shown to modulate RhoA/ROCK activity and thus actin dynamics^20^ (fig 4a). We therefore investigated the contribution of PI3K and its downstream effectors (i.e., AKT, SRC and PYK2) in leukocyte migration under confinement. The treatment of control T cells with specific chemical inhibitor for PI3K increased cell speed, recapitulating the phenotype observed in TTC7A-deficient cells (fig 4e). Similarly, AKT, SRC and PYK2 inhibition in control T cells also increased cell speed (fig 4f-i). In agreement with the hypothesis of reduced PI4KIIIα/PI3K signaling as the cause of altered actin dynamics in TTC7A-deficient cells, exogenous activation of AKT by SC79 in TTC7A-deficient T cells restored cell speed to similar levels as control cells (fig 4j). To further characterize the impact of TTC7A-deficiency in AKT activity, we compared the phosphorylation levels of AKT (pAKT) in HD and TTC7A deficient cells while migrating under confinement. We observed lower pAKT in TTC7A-deficient T cells compared to control, and PI4P supplementation increased pAKT levels of TTC7A-deficient cells (fig 4k-l). These results demonstrate that TTC7A deficiency disrupts the activity of the PI4KIIIα/PI3K/AKT/SRC axis and alters leukocytes motility, highlighting the critical regulatory role of this pathway in actin dynamics and leukocyte migration under confinement.

### Impairment of PI4KIIIα/DIAPH1/actin function in TTC7A-deficient cells disrupts the capacity to migrate through micrometric pores and irregular confined microenvironments

In vivo, leukocyte migration occurs in highly complex microenvironments requiring a high degree of cell deformability^21^. Amongst the different intracellular organelles, nuclear deformation has been shown to be the most important physical limitation to fast interstitial migration of leukocytes^22^. In this context, murine iDC rely on a Arp2/3-dependent actin polymerization to compress and deform the nucleus, which allows fast migration speed in confined 3D microenvironments^23^. To characterize the impact of TTC7A deficiency in the capacity of leukocytes to deform their nucleus, we studied the behavior of control and TTC7A-decifient T cells while migrating through 8 µm microchannels carrying constrictions of 1.5 µm width and 15 µm lenght (fig 5a). In agreement with previous experiments, T cells spontaneously migrated in 8 µm channels devoid of constrictions, while TTC7A-deficient T cells were faster than control T cells (supplementary fig S2c). As previously shown, the presence of 1.5 µm constrictions strongly reduced cell speed of migrating cells ^23^. In the case of control T cells, around 45% of cells were able to deform their nucleus and pass through the constriction (fig 5b and supplementary fig S2d). In contrast, only 25% of TTC7A-deficient T cells were able to pass through 1.5µm constrictions (fig 5b). Of note, the few TTC7A-deficient cells passing the constriction deformed the nucleus likewise control cells (supplementary fig S2d). Next, we aimed to determine whether the inability to pass the constrictions was related to a defective nuclear deformation. To do so, nuclear deformation was quantified in the faction of cells that did not pass the constriction. We defined four different situations depending on the intensity of nuclear deformation: First, cells that did not deform their nucleus at all (group I), cells in which the nucleus entered less than 33% of the constriction (group II), between 33 and 66% of the constriction (group III), and 100% of the constriction (group IV) (fig 5c). Blocked control T cells had an equal repartition among the four different groups (fig 5d-e), suggesting that cell blockage was not determined by impaired capacity to deform the nucleus. In contrast, a large majority of TTC7A-deficient T cells failed to deform the nucleus at the constriction (group I) (fig 5d-e). These results suggest that reduced capacity of TTC7A-deficient T cells to pass through micrometric spaces was caused by an impaired capacity to deform their nucleus.

**Figure 5.**
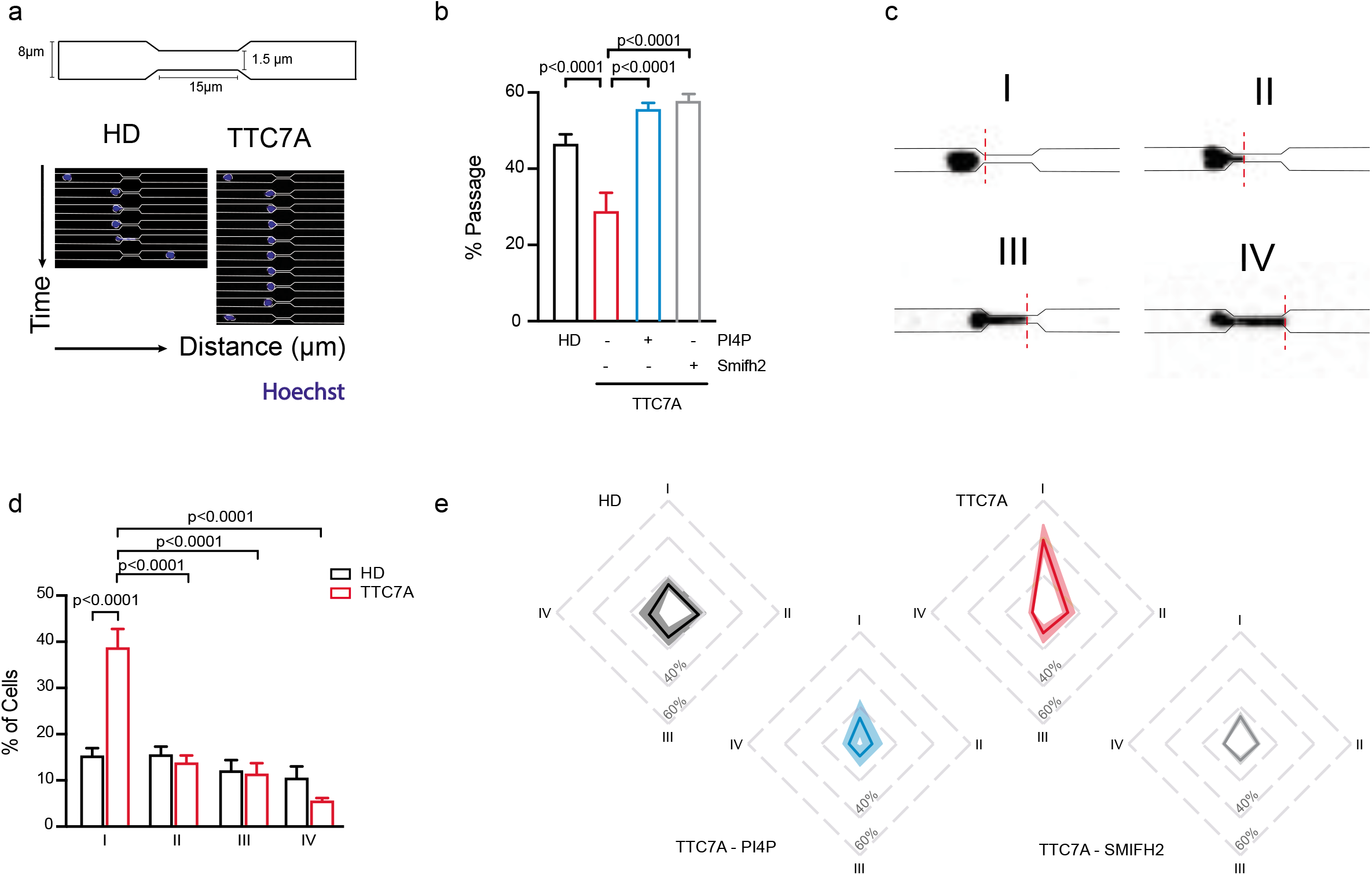
TTC7A is necessary for T cell blast to pass through constrictions: (a) Schematic representation of the migration assay through constricted microchannels (top panel). Lower panel depicts HD and TTC7A-deficient cells passing through a constriction of 1.5µm width and 15 µm of length. Height is fixed at 5 µm in both channels and constrictions. (b) Percentage of HD and TTC7A T cell passing through constrictions of 1.5µm, respectively to the total number of cells reaching the constricted site. TTC7A cells were treated with 25 µM of PI4P (Blue-empty bar) or with 2µM of smifh2 (Gray-empty bar) as indicated. Result from five independent experiments, ≥60 cells analyzed in each condition and experiment. The proportion of passing cells follows a binomial distribution and the p-values were calculated with the chi-square method for each experiment. (c) Classification of the nucleus behavior in non-passing cells. I: no deformation, II: deformation until 33% of the constriction, III: deformation between 33 and 66% of the constriction, IV: deformation covering all (100%) the constriction. (d) Level of nuclear deformation in non-passing cells following groups established in c, respectively to the total number of cells for HD (black-empty bars) and TTC7A (red-empty bars) T cell blast. One-way ANOVA was used to evaluate statistical significance. (e) Distribution of nuclear deformation of non-passing cells respectively to the total number of cells for HD (Black), TTC7A (Red), TTC7A-PI4P (Blue) and TTC7A-Smifh2 (Grey) T cells. Line depicts the mean value and the shadow the standard deviation, 4 independent experiments.

To determine whether the reduced passage rate of TTC7A-deficient cells in 1.5µm constrictions was caused by alterations in the PI4KIIIα signaling pathway, we supplemented TTC7A-deficient cells with PI4P. In this condition, we observed that TTC7A-deficient cells restored their capacity to migrate through 1.5µm constrictions (fig 5b), indicating that the altered actin dynamics characterizing TTC7A-deficient cells impaired the capacity to pass through micrometric pores. Consistently, DIAPH1 inhibition in TTC7A-deficient T cells also restored the capacity of passing through 1.5µm constrictions (fig 5b). In addition, PI4P and Smifh2 treatments re-established nuclear deformability of non-passing TTC7A-deficient T cells to a level comparable to control cells (fig 5e).

To evaluate the impact of TTC7A deficiency in leukocyte migration in a highly complex microenvironment, we assessed cell motility in dense 3D collagen gels. Analysis of random trajectories showed that T cells from TTC7A-deficient patients presented with an overall reduced displacement and a lower capacity to explore their microenvironment, as indicated by a lower mean squared displacement (MSD) compared to control cells (fig 6a-b). To further characterize lymphocyte motility, control and TTC7A-deficient T cells were challenged with the chemokine CCL21. Both cell types responded equally to the chemokine, increasing their directionality to a similar level (fig 6c). Further analysis of cell trajectories during the stable phase of chemotaxis showed equivalent orientation to the chemokine source in both conditions (fig 6d). However, despite a similar directionality, TTC7A-deficient T cells showed slower speed compared to controls (fig 6e). Reduced speed of TTC7A-deficient cells was also observed in random migration, but speed differences between control and TTC7A-deficient cells was increased during chemotaxis (fig 6e).

**Figure 6.**
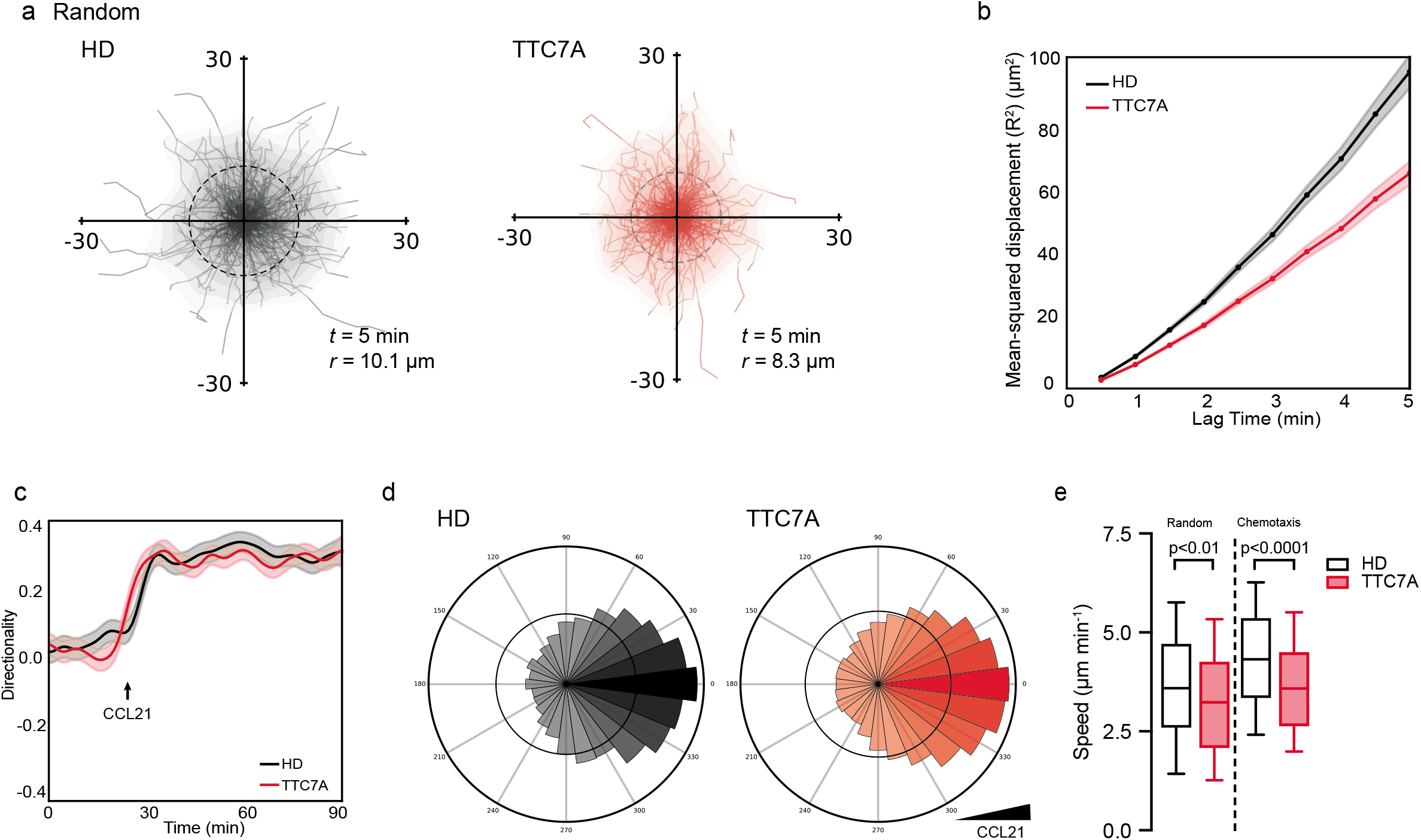
TTC7A is needed to optimize leukocytes motility in highly dense 3D collagen gels: HD (black) and TTC7A (red) T cell blast were embedded into a 3D collagen matrix (3.6 mg/ml) and recorded while migrating in random motion and during chemotactic response towards CCL21 (200 ng/ml). (a) Cell trajectories of HD and TTC7A T cells during random motility. *t* indicates the period of displayed tracking and *r* indicates the mean displacement ratio during that period (b) Mean-squared displacement of T cells migrating randomly in a collagen matrix. Plot corresponds to the analysis of trajectories shown in A. (c) Directionality over time of HD and TTC7A T cells before and after CCL21 addition at 30 min (indicated with an arrow). (d) Directionality of T cells migrating in a collagen gel after addition of CCL21. (e) Mean cell speed of HD and TTC7A cells before (left panel) and after (right panel) stimulation with CCL21. Boxes include the 80% of the points and points represent the higher and lower 10% of points. Mann-Whitney was used to evaluate statistical significance.

These results indicate that actin dynamics’ imbalance as observed in TTC7A-deficient T cells results in a decreased capacity to migrate in 3D landscapes, particularly during chemotaxis.

### TTC7A is essential for preservation of genome integrity and cell survival when migrating in complex 3D microenvironments

Cell migration in confined microenvironments is known to be associated with transient ruptures of nuclear envelope and induction of DNA damage^23, 24^. Therefore, alterations in the capacity of leukocytes to deform their nucleus and migrate in complex confined microenvironments could have deleterious consequences in genomic stability and cell survival. Therefore, it is tempting to propose that impaired migration of TTC7A-deficient T cells could impact cell viability during migration in complex landscapes (fig 7a). To address this, we compared cell viability of control and TTC7A-deficient T cells upon 48 hours of random migration in dense collagen gels, or in absence of confinement (i.e. cell suspension). Cell viability was equivalent between control and TTC7A-deficient T cells in both settings (Fig 7b-c). Next, T cells were submitted to a gradient of CCL21, a condition in which cell speed is profoundly modified (Fig. 6e). In this situation a significant increase in cell mortality of TTC7A-deficient T cells was observed as compared to controls (Fig 7b-c). Then, we sought to evaluate whether the increased mortality of TTC7A-deficient cells was related to DNA damage. To do so, 53BP1 foci staining on nucleus was used as indicator of DNA damage. In keeping with survival assays, we found low levels of DNA damage in absence of confinement or during random migration in both cell types (Fig 7d-e). An increased DNA damage of TTC7A-deficient T cells was observed during chemotaxis (fig 7d-e). These results indicate that impairment in the chemokinetic capacity of TTC7A-deficient T cells in 3D collagen gels is associated with increased DNA damage and reduced cell survival. Strikingly, supplementing TTC7A-deficient T cells with PI4P protected cells from DNA damage and death during directional migration in collagen gels. These results indicate that alterations in actin dynamics caused by defective PI4KIIIα/PI3K/AKT/RHOA signaling in TTC7A-deficient cells are associated with an impaired nuclear deformation, resulting in increased DNA damage and reduced survival during directional migration in complex 3D microenvironments (fig 7f).

**Figure 7.**
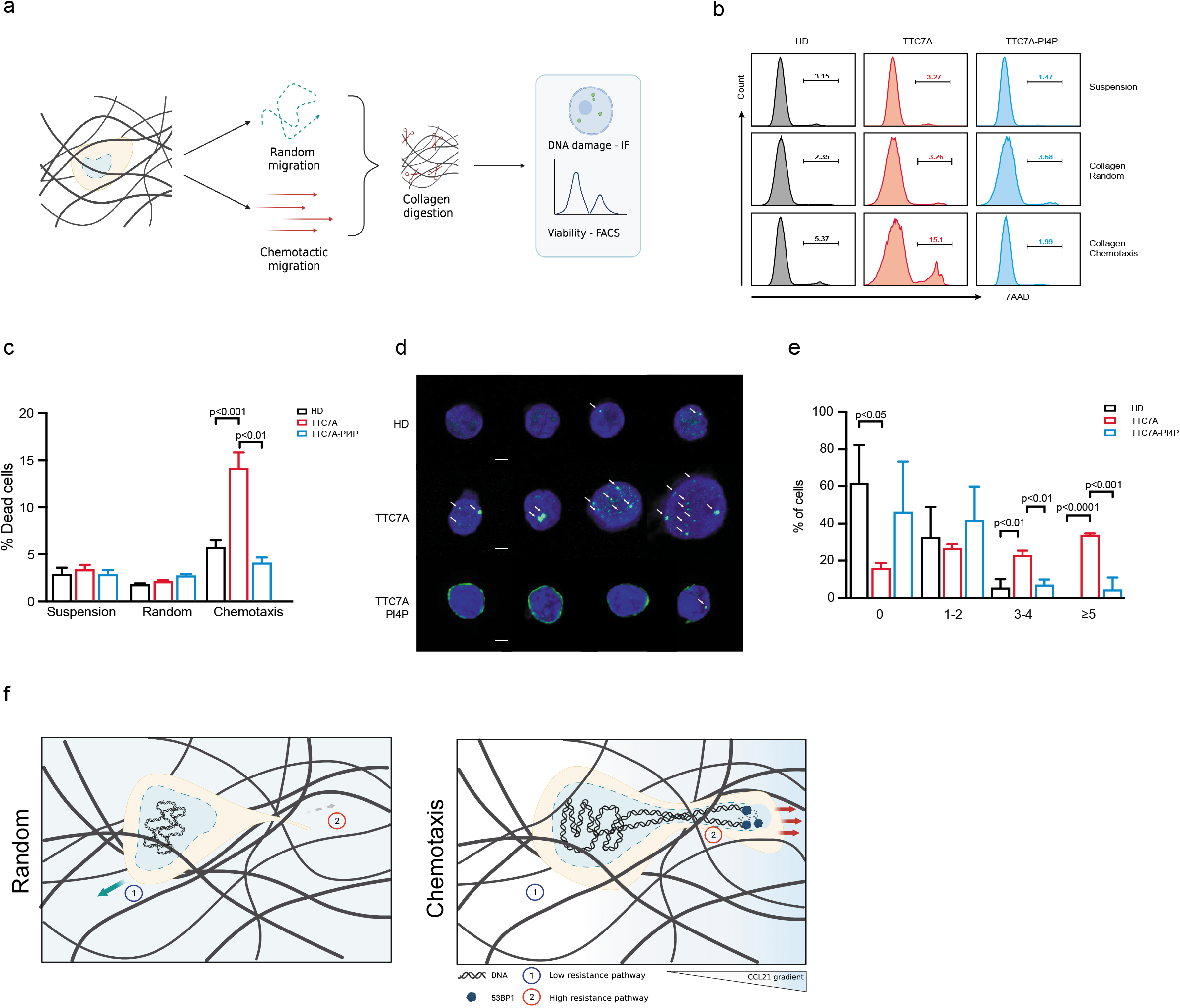
TTC7A is required for T cell blast survival when migrating in highly dense microenvironments: (a) Experimental design, T cell blast were embedded into highly dense collagen (3.6 mg/ml), allowed to migrate randomly or in response to a chemotactic gradient of CCL21 (200 ng/ml) during 48h, collagen was digested with collagenase (1mg/ml in 2.5 mM CaCl_2_ for 1 h) and recovered cell were used to evaluate DNA damage and viability. (b) Representative histograms depicting the percentage of 7AAD positive cells in suspension (Top panels), random collagen (Middle panels) and chemotactic collagen (Bottom panels) for HD (Black), TTC7A (Red) and TTC7A-PI4P (Blue) T cell blast. (c) Percentage of dead cells for T cell blast in suspension and cells recovered from random and chemotactic collagen. HD (Black-empty bars), TTC7A (Red-empty bars) and TTC7A-PI4P (25 µM, blue-empty bars). Result including 8 independent experiments. Bars represent mean and error bars SEM. One-way ANOVA test was applied for statistical analysis. (d) 53bp1 staining of HD (Top panel), TTC7A (Middle panel) and TTC7A-PI4P (Bottom panel) T cells recovered from collagen. Scale bar 3µm. (e) Quantification of the number of 53bp1 foci per cell for HD (n=117), TTC7A deficient (n=111) and TTC7A deficient T cells treated with PI4P (25µM of n=70). Bars represent mean and error bars SEM. One-way ANOVA was used to evaluate statistical significance. (f) Schematic representation of T cells while migrating randomly in collagen (Top panel) and in response to a chemotactic gradient (Bottom panel).

These results highlight TTC7A as a critical regulator of leukocyte migration and cell survival under confinement and emphasize that perturbations of this process may contribute to dysregulated immune homeostasis observed in TTC7A-deficient patients.

## Discussion

To date, TTC7A has been shown to participate in several cellular functions ^2, 10, 11, 25, 26^. However, it is not clear how alterations in these processes contribute to the immune phenotype characterizing TTC7A-deficient patients. The present report shows that TTC7A is a regulator of leukocyte migration under confinement (e.g. the interstitial space) in human and mice. TTC7A controls F-actin polymerization by ensuring the activity of the PI4KIIIα/PI3K/AKT/RHOA regulatory axis (fig 8). The synthesis of PI4P is the first step of a signaling pathway leading to PI(3,4,5)P_3_ production, AKT activation, and eventually RHOA activity ^20^, among many others ^27^. In TTC7A-deficient cells, the defective kinase activity of PI4KIIIα decreases the PI4P pool ^8, 28^ and leads to RHOA hyperactivation. Subsequently, RHOA destabilizes the autoinhibited conformation of the DIAPH1, promoting the polymerization of actin filaments and persistent motility ^29, 30^. It is known that the coordinated action of mDia1 and Arp2/3 controls murine DC migration by promoting actin polymerization in different subcellular locations ^14^. Our results show that a similar mechanism occurs in human leukocytes. The increase DIAPH1 activity observed in TTC7A-deficient cells suggests that a competition between DIAPH1 and ARP2/3 for free actin monomers could modulate their activities, and thus actin dynamics and cell migration. Such regulatory loop has been described in yeasts, amoebas, drosophila and mammalian cells, and depletion or inhibition of one actin regulator increases the activity of the other ^31 32^.

**Figure 8.**
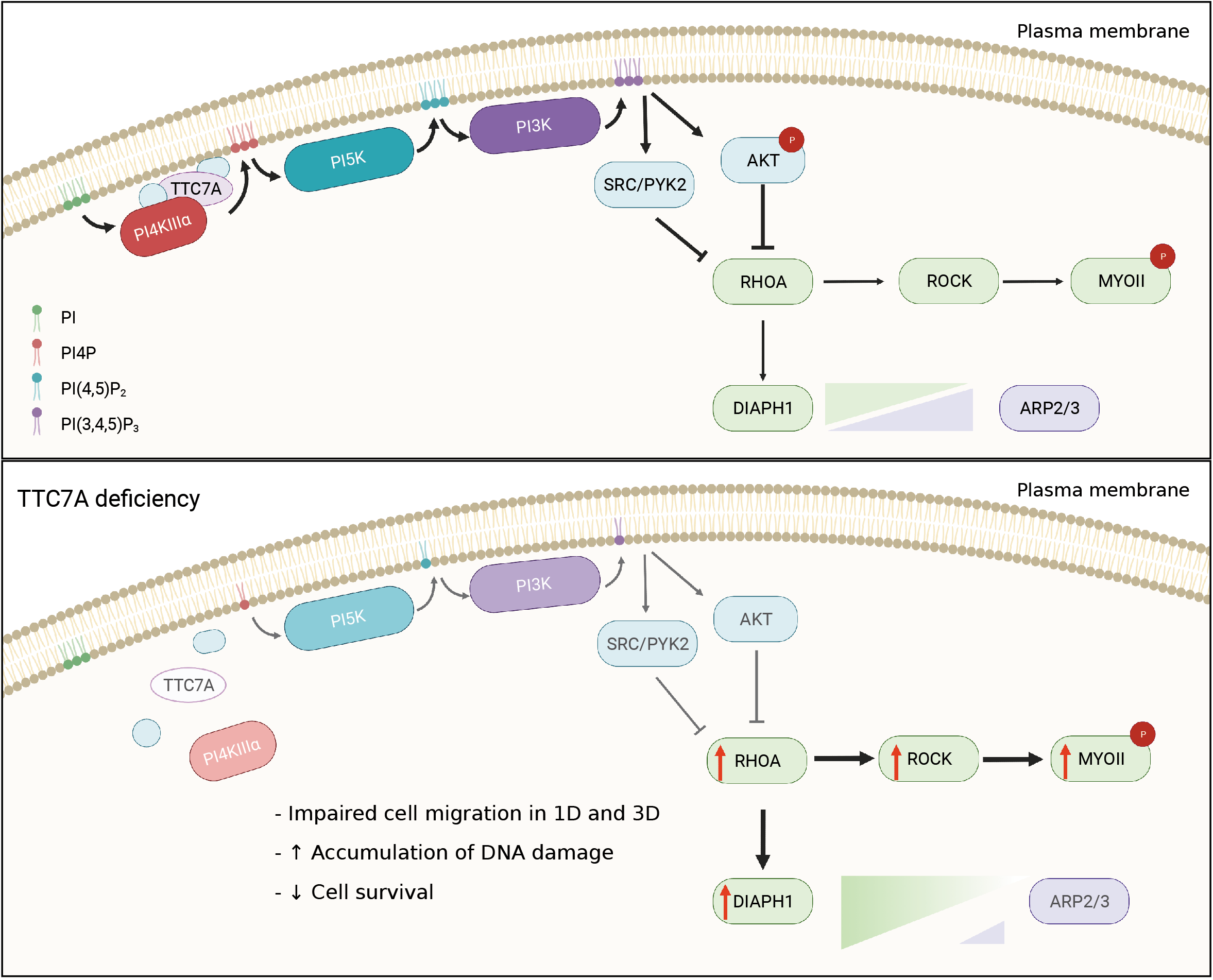
Signaling pathway controlled by TTC7A. Schematic representation of TTC7A/PI4KIIIα pathway and its role in the control of the actin cytoskeleton dynamics in homeostatic conditions (Top panel) and in TTC7A deficiency (Bottom panel).

Surprisingly, it was found that the impact of TTC7A deficiency on leukocyte migration was dependent on the complexity of the surrounding microenvironment, since TTC7A-deficient cells migrated faster in 1D microchannels but were less efficient to migrate through microenvironments requiring a high degree of deformation. This is in agreement with data showing that leukocyte motility in irregular confined microenvironments depends on the ability of the cells to deform ^33, 34^. During interstitial migration, murine leukocytes deform their nucleus in an ARP2/3-dependent process ^22, 23^. As a consequence of this deformation, the nucleus suffers a high degree of mechanical stress, which can be associated with breaks in the nuclear envelope and exposure of nuclear DNA to the cytoplasm ^35–37^. In control cells, migration-induced DNA damage is prevented by the fast resealing of the nuclear envelope ^36^. In context of TTC7A deficiency, alterations in actin dynamics reduce the nuclear deformation capacity, leading to accumulation of DNA damage, and reduced cell survival during chemotactic response in confined complex microenvironments. The deleterious effect of TTC7A deficiency in cell survival was only seen in response to chemokines. This result indicates that during random migration, cells can choose a trajectory imposing the lowest degree of nuclear deformation, which is in agreement with data showing that nuclear deformation is the limiting factor for leukocyte migration ^24, 38^. Our results bring forward the possibility that alterations in leukocyte migration could constitute a pathophysiological mechanism underlying the progressive lymphopenia reported in TTC7A-deficient patients (fig 8).

It is worth noting that TTC7A is a nuclear component regulating chromatin organization and gene expression ^10^. Moreover, the nucleus is a mechanosensitive organelle able to perceive cell compression ^39, 40^, thus further studies should address whether TTC7A could also indirectly affect leukocyte migration by modifying nuclear stiffness. In particular, whether or not TTC7A impacts on the nuclear envelop, known to limit nuclear deformation and immune cell migration warrants further analysis ^23, 41^.

Our work raises intriguing questions as to whether alterations in leukocyte migration contribute to immunodeficiency in other types of PIDs, and whether mechanisms are comparable to TTC7A deficiency. It has been shown that mutations in DOCK8 lead to leukocyte susceptibility to undergo a form of cell death known as cytothripsis ^42^. Cytothripsis is caused by the loss of front/rear coordination during displacement leading to the mechanical break of the cell ^42^. Notably, cytothripsis was not observed in TTC7A-deficient leukocytes, suggesting that TTC7A is not required to preserve cell shape during motility. Leukocyte migration can also be defective in other PID associated with defects in the production of PI4P (or other metabolites of this pathway), as it is the case for the recently described mutations in *PI4KA* ^43, 44^ in patients with a clinical phenotype partially resembling MIA-CID patients ^1, 45^ (i.e. caused by *TTC7A* null mutations). Based on our data, leukocytes from *PI4KA*-deficient patients will likely display similar motility defects, with reduced survival when migrating in dense microenvironments. Alterations in leukocyte migration could also contribute to pathophysiology in other conditions such as defects of DNA damage response machinery ^46, 47^. Indeed, ATR and ATM are required for repair of DNA damage and preservation of DNA integrity during migration under confinement ^35, 36^. However, it is not known whether alterations in this process contribute to clinical phenotypes.

In conclusion, we unveil TTC7A as a critical regulator of actin dynamics, as it allows leukocytes to efficiently migrate under confinement. We show that the reduced capacity to deform their nucleus impairs interstitial migration of TTC7A-deficient leukocytes, leading to an accumulation of DNA damage, and reducing cell survival. Moreover, we propose that alterations in nuclear mechanics in leukocytes during confined migration could be a previously unappreciated pathophysiological mechanisms contributing to the development of progressive leukopenia and immunodeficiency.

## Acknowledgments

The authors would like to thank to the Imagerie Cellulaire and the Bio-Image Analysis Platforms of the SFR Necker, for their technical support with the microscopy experiments and image analysis. Figures 7a, 7f and 8 were created with biorender.com. Lifeact-GFP mice were kindly donated by M Sixt (Austria).

TG was supported by the International PhD program of the Imagine Institute and the Fondation Bettencourt Schueller. ML was supported by the Imagine Institute (WP05T012). MB was supported by ATEurope and IPGG HRHG.

This work was supported by the State funding from the Agence Nationale de la Recherche under “Investissements d’avenir” program (ANR-10-IAHU-01), and the French National Institute of Health and Medical Research (INSERM). The laboratory of FE Sepulveda receives funding and supports from the Agence Nationale de la Recherche (ANR-18-CE15-0017), the ARC foundation (PJA 20191209614), La Ligue Contre le Cancer (RS19/75-79, RS20/75-30), and the Imagine Foundation. GM was funded by La Ligue Contre le Cancer (RS21/75-3) and the ARC foundation (PJA 20181207755). PV was funded by the Agence Nationale de la Recherche (ANR-16-CE13-0009), the Emergences Canceropole (SYNTEC project) and Labex-IPGG (ANR-10-IDEX-0001-02 PSL and ANR-10-LABX-31).

## Authors contributions

TG conceived and performed experiments, analyzed data, and wrote the manuscript. ML, MB, CL, MTED, MK, GLL performed experiments and analyzed data. DM, BN, AF contributed biological material and discussed data. AF, GM, GSB analyzed data, contributed ideas, and edited the manuscript. PV conceived and performed experiments, analyzed data, and wrote the manuscript. FES conceived the study, analyzed data, wrote the manuscript and supervised the overall research. All authors revised, edited and approved the final version of the manuscript.

## Methods

### Patients

TTC7A-deficient patients have been previously reported^2, 4^, all of whom had given their consent to participation in the study. Patients with the following biallelic mutations in the *TTC7A* gene were included: L304fsX59, E71K, R325Q. A ficoll-paque density gradient was performed to recover the peripheral blood mononuclear cells (PBMC) from patients and healthy donors. Cells were activated with 5µg/ml phytohemagglutinin (PHA, Sigma) and 100 U/ml of IL-2 (PeproTech) for 3 days and cultured in RPMI 1640 medium supplemented with 10% of FBS and 1% penicillin/streptomycin (all Gibco, Thermofisher). To obtain monocytes, CD14+ cells were purified using CD14 positive selection kit (BD Bioscience) and used immediately for migration experiments. LCL were generated from patients’ PBMC immortalized by infection with Epstein-Bar Virus as previously described ^1^.

### Mice

Heterozygous Balb/cByJ fsn (CByJ.A-Ttc7fsn/J) mice were obtained from the Jackson Laboratory and Lifeact-GFP (C57bl/6) were donated from M.Sixt (IST, Austria)^16^. In order to work with reporter *fsn*-LifeAct mice we bred both lines (CB6F1) and performed all experiments with the F1 generation of this breeding. All mice were maintained in SPF (Specific Pathogen Free) facility and handled according to national and institutional guidelines.

### Mouse-derived cells

To obtain bone marrow derived dendritic cells, bone marrow from the femurs of *fsn* and littermate control mouse (3 to 4 weeks old) was recovered and cultured during 10 days in Iscove’s modified dulbecco’s medium (IMDM) supplemented with 10% FBS, 1% penicillin/streptomycin, 50µM β-mercaptoethanol and supernatant containing 50ng/ml of granulocyte-macrophage colony stimulating factor (GM-CSF). Semi-adherent cells were recovered and either used as immature DC (iDC) or treated with 100 ng/ml of LPS (mDC) for 30 min. For microchannels migration experiments cells were stained with 200 ng/ml of Hoechst 33342 (Molecular Probes) for 30 min and resuspended at 20×10^6^cells/ml, acquisition took place between 4 and 16 h after activation. For collagen migration cells were cultured overnight after activation to reach full maturation and analyzed between 24 and 36 h post-treatment.

T cells were recovered from the spleen by perfusion with PBS, red blood cell lysis was done using RBC lysis buffer (Biolegend) during 1 minute in ice. Cells were activated in a 24-well plate coated with anti-CD3 (clon: 500A2, Biolegend) and soluble anti-CD28 (clon: 37.51, Biolegend), for 3 days.

### Micro-devices

Microdevices were prepared as previously described ^48^. Briefly, devices were fabricated using polydimethylsolixane (PDMS) and custom-made molds, coated with 10ug/ml fibronectin from bovine plasma (Sigma) for 1 h. In the case treatments, devices were incubated with media containing the drug for at least 30 min at 37°C. Migration was recorded using a Zeiss Axio Observer Z1 (Hamamatsu digital camera C11440, Carl Zeiss) microscope, with a time-lapse of 2 min using a 10X (N.A. 0.45) dry objective. The image analysis was done using ImageJ (NIH). In short, kymographs for each channel were generated using a semi-automatic macro and single cell trajectories were manually isolated. Trajectories were then analyzed with a custom-made Matlab (Mathworks) code to determine the cell speed. For the analysis of the constrictions same macro was used for the generation of the kymographs and a second semi-automated macro was used for calculating the percentage of cells passing through the constriction. The percentage corresponds to the number of cells that pass respectively to the total number of cells reaching the constriction.

### Internalization assay

Cells were allowed to migrate overnight in a close microchannels system, after migration the media was removed and replaced with 40KDa-Dextran coupled to Texas red. Acquisition of images was done during the first hour after the exchange of media, using Zeiss Axio Observer Z1 (Hamamatsu digital camera C11440, Carl Zeiss) microscope and an oil-40X objective (N.A. 1.4)

### Migration in collagen gels

The migration of leukocytes in collagen gels was performed as described previously ^21^. Briefly, cells at a concentration of 2×10^6^/ml were mixed with rat-tail collagen type 1 (Corning) to a final concentration of 3.6 mg/ml in a basic pH environment and loaded into a PDMS custom made chamber and allowed to polymerize at 37°C for 15 min. Acquisition was performed overnight (37°C and 5%CO_2_) using a DMi8 inverted microscope (Leica) and a ×10 dry objective (NA 0.40 phase), with a timelapse of 1 minute. After 30 min of recording, CCL21 (Peprotech) was added to the media at a final concentration of 200 ng/ml to form a chemotactic gradient. The recovered video was analyzed using ImageJ, an average subtraction and mean filters were applied to the videos in order to clean the images, which later were tracked using Imaris (Bitplane). The tracking data was then analyzed using custom-made software to extract the relevant kinetic information.

### Antibodies and drugs

For FACS analysis the following antibodies were used: anti-CD11c APC and anti-I-A/I-E APC-Cy7 (Biolegend), anti-CD40 PeCy7, anti-CD80 Fitc, anti-CD86 Pacific Blue (Sony), anti-CCR7 PE (Macs), anti-IgG2aκ PE (BD Pharmigen), for mouse. Anti-CD3 Percp (Macs), anti-CD4 Pacific Blue, anti-IgG2aκ Pecy7 (Biolegend), anti-CD8 PE, anti-CCR7 Pecy7 (BD Pharmigen) for human.

For Western-blot and immunofluorescence: GAPDH (clon: 6C5, Merck), mDia1 (clon: 51/mDia1, BD Bioscience), ARP2 (polyclonal, Abcam), phospho-akt AlexaFluor647 (Ser473, clon: 193H12, Cell signaling). Flow Cytometry acquisition was done in a Gallios flow cytometer (Beckman Coulter) and analyzed with the software Flowjo (Treestar).

The drugs and concentrations used in this study are the following Blebbistatin (50µM), Y-27632 (7.5 - 10 µM), Smifh2 (2µM), Calpeptin (30 mg/ml), CK666 (25µM), GSK-A1 (100nM), all from Sigma. BF738735 (1.7 µM), Tenasilib (45 nM), MK2206-2HCl (7.5 nM), SC79 (16.5 µM), Src Inhibitor1 (1µM), PP2 (10µM), Defactinib-HCl (1µM), PF-562271 (10µM) from Medchemexpress. Pertusis-toxin (200 ng/ml, Thermofisher). PI4P was acquired from Echelon Bioscience and used as instructed, briefly carriers and lipids were resuspended in aqueous buffer (15mM NaCl, 4mM KCl, 20 mM Hepes), mixed at equal concentration and incubated at RT for 10 min to allow the complexes to form, dilute to appropriate concentration and incubate with cells during 30 min at 37°C.

### Enzyme-Linked Immuno Sorbent Assay

50×10^3^ iDCs were plated in P96 round bottom and activated overnight with increasing concentrations of LPS (0, 0.4, 2, 10 and 50 ng/ml). Supernantant was recovered and used for measuring IL-6 and TNF-alpha (ebioscience) secretion by ELISA according to manufacturer’s protocol. Absorbance was read in 2030-040 Victor X4 (Perkin Elmer) at 450nm.

### Immunofluorescence

DCs were allowed to migrate in 8µm width microchannels for 16 h and then fixed with 4% PFA in PBS for 45 min at RT. The PDMS block was removed from the coverslip containing the cells that migrated fixed in a polarized state. The cells in the coverslip were permeabilized for 2 min with PBS+0.1% of Triton X-100 at RT and blocked for 1 h with PBS + 2% BSA also at RT. Primary antibody incubation was done overnight at 4°C in the dark, together with phalloidin Flash Green (Biolegend) and Hoechst 33342. Then coverslips were incubated for 45 min with the proper secondary antibody at RT. Washed and mounted using proGold anti-bleaching solution (Thermofisher).

The analysis of immunofluorescence images was done using ImageJ as described before^14^.

Shortly, cells were aligned, normalized by size and the average fluorescence intensity of the population was determined for each staining of interest (supplementary fig S3). The values of the intensity of fluorescence were determined with a mask of the back and front of the cell, excluding the nucleus using Icy.

### Western blot

DCs were lysed for 10 min in ice with a buffer containing 0,1% NP40, 50mM Hepes, 150 mM NaCl, 1mM MgCl2 and 10% glycerol, supplemented with anti-protease and anti-phosphatase tablet (Roche). 20µg of each sample was loaded into a 4-12% Bis-Tris Gel (Invitrogen) and transfer into a nitrocellulose membrane. After blocking and antibody incubation the membrane was revealed using ECL-Plus western blotting substrate (Pierce) and read using the Chemidoc imaging system and Image lab (BioRad) was used to analyze the blots.

### Viability in collagen

High concentration collagen gels were prepared by mixing rat tail collagen type I (Corning, 3.6 mg/ml) together with T cell blast at 4°C and a basic pH, seeded into a 48 well plate and allowed to polymerize for 13 min at 37 °C. Media containing 200 ng/ml of recombinant CCL21 (Peprotech) was added to the well to generate a chemotactic gradient. After 48h a solution of Collagenase type I (Worthington, 1mg/ml) in CaCl_2_ (2.5 mM) was added for 1 h at 37°C for digestion of the collagen. Cells were stained with anti-CD3 Percp, anti-CD4 Pacific blue (Macs and Biolegend, respectively) and 7AAD was used as viability marker (Biolegend) to quantify cell death by FACS. Also, recovered cells were used for immunofluorescence against 53BP1 as previously described. Briefly, cells were allowed to attach to cover slides overnight and then fixed with 4% PFA for 20 min, treated with 0.1 M of Glycine for 30 min, permeabilized with 0,5% Triton X100, blocked during 1 h with PBS + 2% BSA. Incubation with 53BP1 antibody (polyclonal, Novus biologicals) for 90 min at RT, washed, incubated with proper secondary antibody for 1 h at RT and mounted using proGold anti-bleaching solution (Thermofisher).

### Statistics

All data analysis was performed with GraphPad Prism 9 for MacOS (GraphPad Software). Data obtained from migration experiments was evaluated for normal distribution using the D’Agostino & Pearson test, and comparisons between conditions were performed using Mann-Whitney/unpair t-test (for two conditions) or one-way ANOVA/Krustal-Wallis test (more than two conditions), depending on the normality result. The passage through microconstrictions follows a binomial distribution and p values were calculated using the chi-squared method for each experiment and set of comparisons. Statistical differences were considered when *p<0.05, **p<0.01, ***p<0.001 and ****p<0.0001.

---

**Supplementary figure 1.**
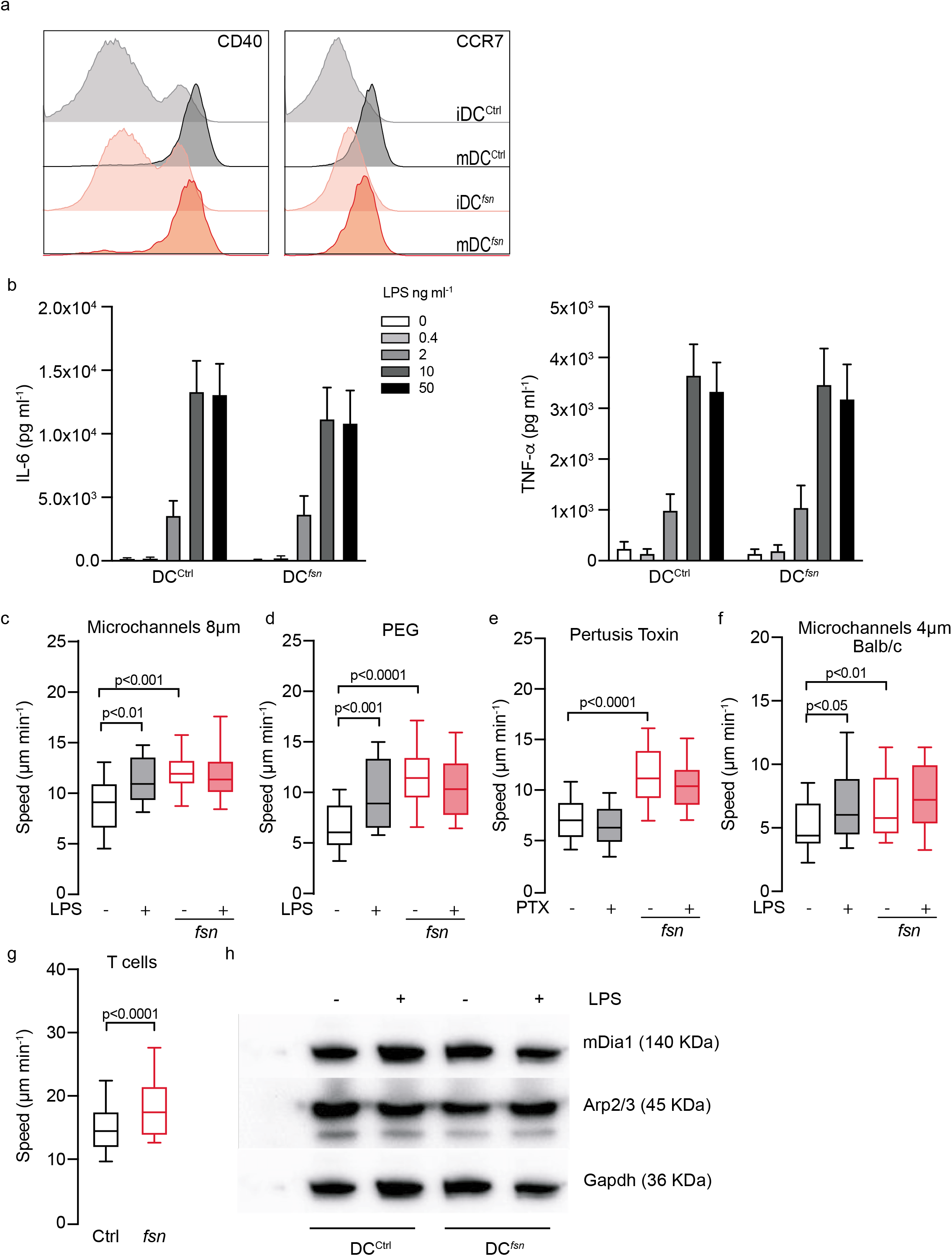
(a) Flow cytometry analysis of CD40 and CCR7 iDC^ctrl^ (Light grey), mDC^ctrl^ (Dark grey), iDC*^fsn^* (Light red) and mDC*^fsn^* (Dark red). (b) Quantification of cytokine secretion from of iDC^ctrl^ and iDC*^fsn^*. Elisa test for IL-6 (left panel) and TNF-a (right panel) measured in pg/ml. 50×10^3^ cells stimulated with indicated concentrations of LPS. (c-g) Mean instantaneous speed of iDC^ctrl^ (Black-empty bars), mDC^ctrl^ (Black-filled bars), iDC*^fsn^* (Red-empty bars) and mDC*^fsn^* (Red-filled bars), under the specified conditions. Boxes include the 80% of the points and bars represent the higher and lower 10% of points. One-way ANOVA test was used to evaluate statistical significance. One representative experiment out of 2 or 3 is presented. (c) DCs migrating in channels of 8µm x 5 µm, fibronectin-coated. (d) DCs migrating in channels of 4 x 5 µm coated with polyethynel glycol. (e) DCs migrating in channels of 4 x 5µm fibronectin-coated, treated or not with 200 ng/ml of pertussis toxin as indicated. (f) DCs from Balb/c background, migrating in channels of 4 x 5 µm, fibronectin-coated (g) T cells from ctrl and *fsn* mice migrating in channels of 4 x 5µm fibronectin-coated. Mann-Whitney test was used to evaluate statistical significance. (h) Immunoblot analysis of mDia1 (140 KDa), Arp2/3 (45 KDa) and GAPDH, from proteins extracted of iDC^ctrl^, mDC^ctrl^ iDC*^fsn^* and mDC*^fsn^*.

**Supplementary figure 2.**
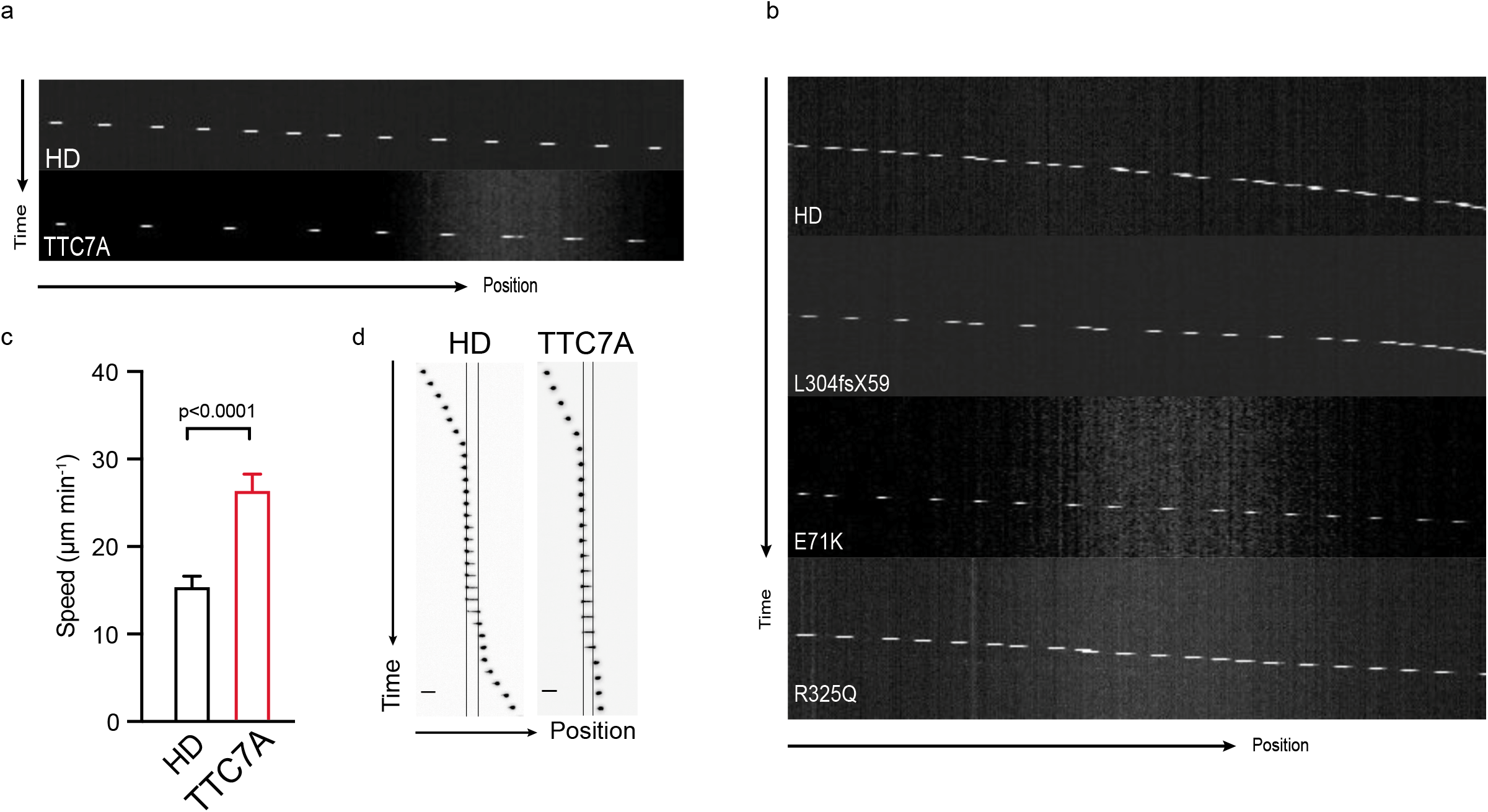
(a) Representative kymographs of primary monocytes in 4µm x 5µm fibronectin-coated microchannels. HD (Top panel) and TTC7A (Bottom panel). (b) Representative kymographs of Lymphoblastoid cell lines (LCL) carrying three different mutations in the *TTC7A* gene as indicated, in 8µm x 5µm fibronectin-coated microchannels. HD (Top panel) and TTC7A (Bottom panels). (c) Mean instantaneous speed of HD (Black-empty bar) and TTC7A deficient (Red-empty bar) T cell migrating in 8µm x 5µm fibronectin-coated microchannels. Bars represent mean and error SEM. Mann-Whitney test was used to evaluate statistical significance. (d) Representative montage of HD and TTC7A T cell blast passing through a 1.5 µm constriction over time. Each slice represents 1 min and bar 15 µm.

**Supplementary figure 3.**
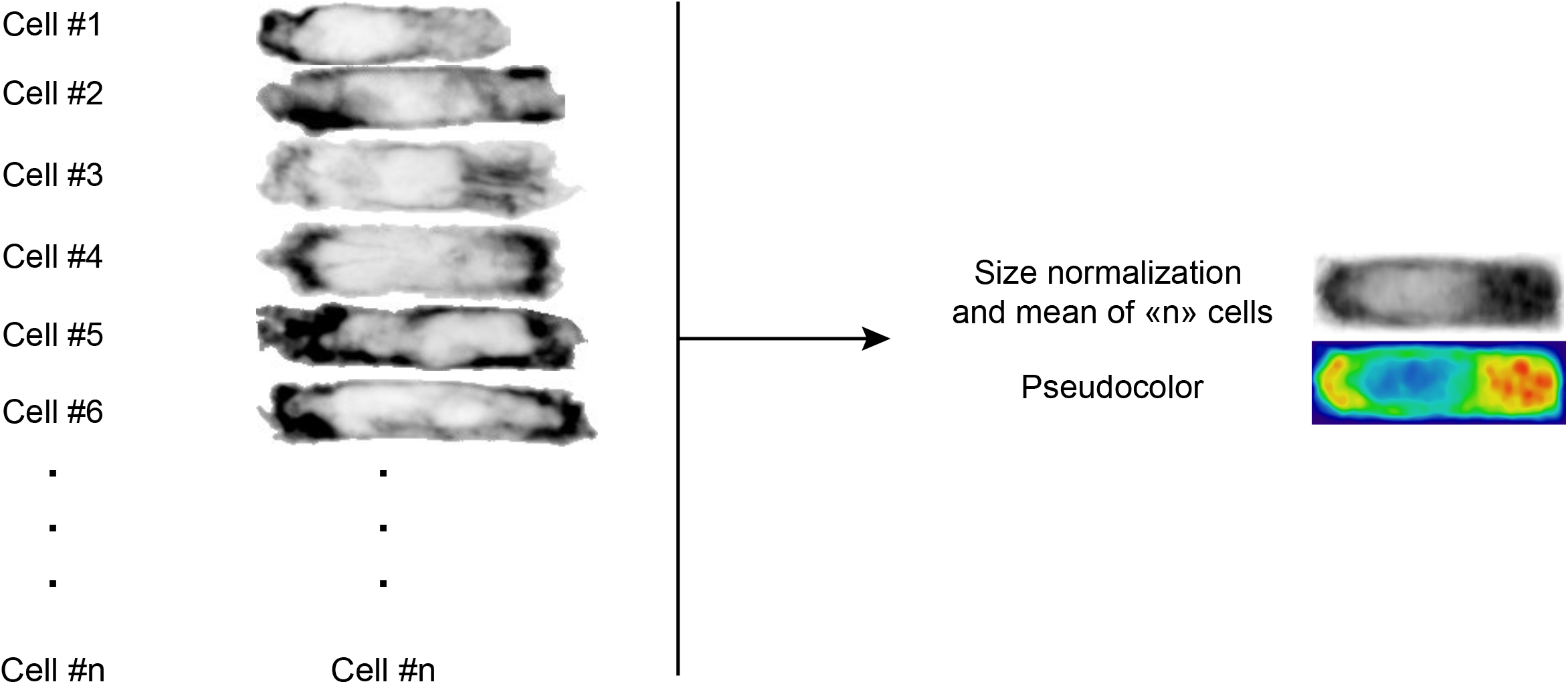
Analysis strategy of immunofluorescence in polarized cells. Using ImageJ, each cell was separated and oriented in the same direction of migration; the contour of the cell was drawn with a phalloidin-actin mask. Cells were then aligned and normalized by size, the mean of “n” cells was obtained and pseudocolor was applied to the result.

